# Genomic Diagnostics for Drug-Resistant Mycobacterium tuberculosis: Computational Prediction of Antimicrobial Resistance

**DOI:** 10.64898/2026.05.25.727578

**Authors:** Mohammadali Serajian, Yunheng Han, Christina Boucher

**Author notes:** Corresponding author: Christina Boucher.

## Abstract

Tuberculosis (TB) remains a leading cause of infectious disease mortality, and the continued emergence of drug-resistant *Mycobacterium tuberculosis* (MTB) strains threatens the effectiveness of standard treatment regimens. Culture-based antibiotic susceptibility testing (AST) remains the clinical reference standard for resistance determination but typically requires six to eight weeks, delaying initiation of optimized therapy for patients with drug-resistant disease. Whole-genome sequencing (WGS)-based approaches provide a rapid alternative for predicting antimicrobial resistance directly from genomic data and are increasingly being incorporated into diagnostic workflows. This survey reviews computational approaches for genomic resistance prediction in MTB, focusing on two major classes of methods: catalog-based tools that identify established resistance-conferring variants, and *de novo* machine learning approaches that infer resistance from genome-wide sequence features. We examine the strengths and limitations of these approaches with respect to interpretability, scalability, computational requirements, and concordance with phenotypic testing. We further discuss emerging directions in quantitative minimum inhibitory concentration (MIC) prediction, challenges in pyrazinamide susceptibility testing, and the limited availability of resistant isolates for newer and repurposed drugs used in multidrug-resistant TB (MDR-TB) and extensively drug-resistant TB (XDR-TB) treatment regimens. Continued expansion of paired phenotypic and genomic datasets, standardized MIC testing protocols, and rigorous lineage-aware evaluation frameworks will be essential for improving the clinical reliability and global deployment of genomic resistance prediction for tuberculosis diagnostics.

## 1 Introduction

Tuberculosis (TB) is a disease known for millennia and is caused by a group of closely related bacteria, collectively known as the Mycobacterium tuberculosis complex (MTBC) [17]. TB is the leading cause of death due to a single infection agent [53], and has led to more than a billion deaths in the past 200 years [32]. This global endemic infection spreads from person to person through airborne transmission [13]. TB is most commonly reported as pulmonary disease but it can also manifest in other forms such as extrapulmonary TB, most frequently involving the lymph nodes and pleura, and less commonly bones, joints and the meninges [13, 33].

Sustained global control efforts have substantially reduced TB mortality, from 2.4 million deaths in 1998 to 1.2 million in 2024, with an estimated 83 million deaths averted between 2000 and 2024 [53, 54]. This progress followed the World Health Organization (WHO) declaration of TB as a global emergency and the establishment of worldwide surveillance in 1995 [37], which addressed earlier gaps in burden estimation amid weakening public health systems and HIV co-infection [36]. However, TB incidence has remained broadly stable over the same period [53].

A key factor underlying this persistent incidence is the continued reliance on legacy tools and a substantial gap between estimated incidence and case notification: approximately 25% of incident cases were not diagnosed in 2024 [53]. Diagnosis and monitoring still rely on microbiological confirmation and microscopy-based methods constrained by the slow-growing, fastidious nature of *Mycobacterium tuberculosis* (MTB), with sensitivity further degraded in paucibacillary disease where bacterial burdens fall below current detection limits [13, 17]. The TB report 2025 demonstrated a treatment success rate of 88% for drug-susceptible TB treated with first-line regimens in 2022, whereas treatment success for multidrug-resistant TB (MDR-TB)/RR-TB was 71% in 2022, albeit a 42% improvement compared to 2012 [54].

Antimicrobial chemotherapy remains the cornerstone of TB control [44]. The recommended treatment relies on multi-drug regimens and can span 4 to 24 months depending on the drug-resistance profile, the initial clinical picture, and the patient’s response over the course of therapy [13]. The impact of therapy is limited due to complexities such as drug toxicity, patient compliance, and emergence of drug resistance [44]. Drug-resistant TB (DR-TB) is classified by the extent of resistance to anti-tuberculosis agents, most importantly rifampicin, which inhibits the MTB RNA polymerase [11]. Rifampicin is considered the most powerful first-line antibiotic capable of shortening the duration of treatment [13, 56]. Additionally, MDR-TB is defined as MTB resistant to both rifampicin and isoniazid [1]. In clinical practice, Rifampicin-resistant TB (RR-TB) is commonly treated as the operational entry point into second-line antibiotic therapy, because RR-TB is managed with regimens aligned with those used for MDR-TB, regardless of concurrent isoniazid susceptibility [13].

Effective TB management depends on the rapid detection of drug resistance and timely initiation of an effective regimen, a priority embedded in the WHO TB control strategy [52]. The clinical stakes are illustrated by a nationwide Rwandan cohort in which reducing the median diagnostic delay from 88 to 1 day and the therapeutic delay from 76 to 3 days between 2006 and 2016 coincided with a decline in rifampicin-resistant TB mortality from 30.8% to 6.9%; a total delay of at least 100 days was independently associated with more than two-fold higher odds of death [30]. With drug-resistant TB estimated to account for 13% of all antimicrobial resistance (AMR)-attributable deaths worldwide [13], scalable and rapid diagnostics are a pressing need.

Treatment selection has traditionally relied on *in vitro* drug-resistance profiling, which remains the reference standard but can take weeks to months depending on the antibiotic [13], limiting the timely regimen selection envisioned by WHO guidance [52, 59]. These constraints have motivated faster alternatives, including molecular diagnostics recommended by WHO for use while culture-based results remain pending, and computational approaches that infer resistance profiles directly from whole-genome sequencing (WGS) or targeted next-generation sequencing data [49, 59]. The latter potentially circumvent the speed and infrastructure limitations of phenotypic testing, and are increasingly informed by genome-wide association studies (GWAS) that identify novel resistance-associated loci and characterize their phenotypic effects [14]. See Figure 2.

**Figure 1:**
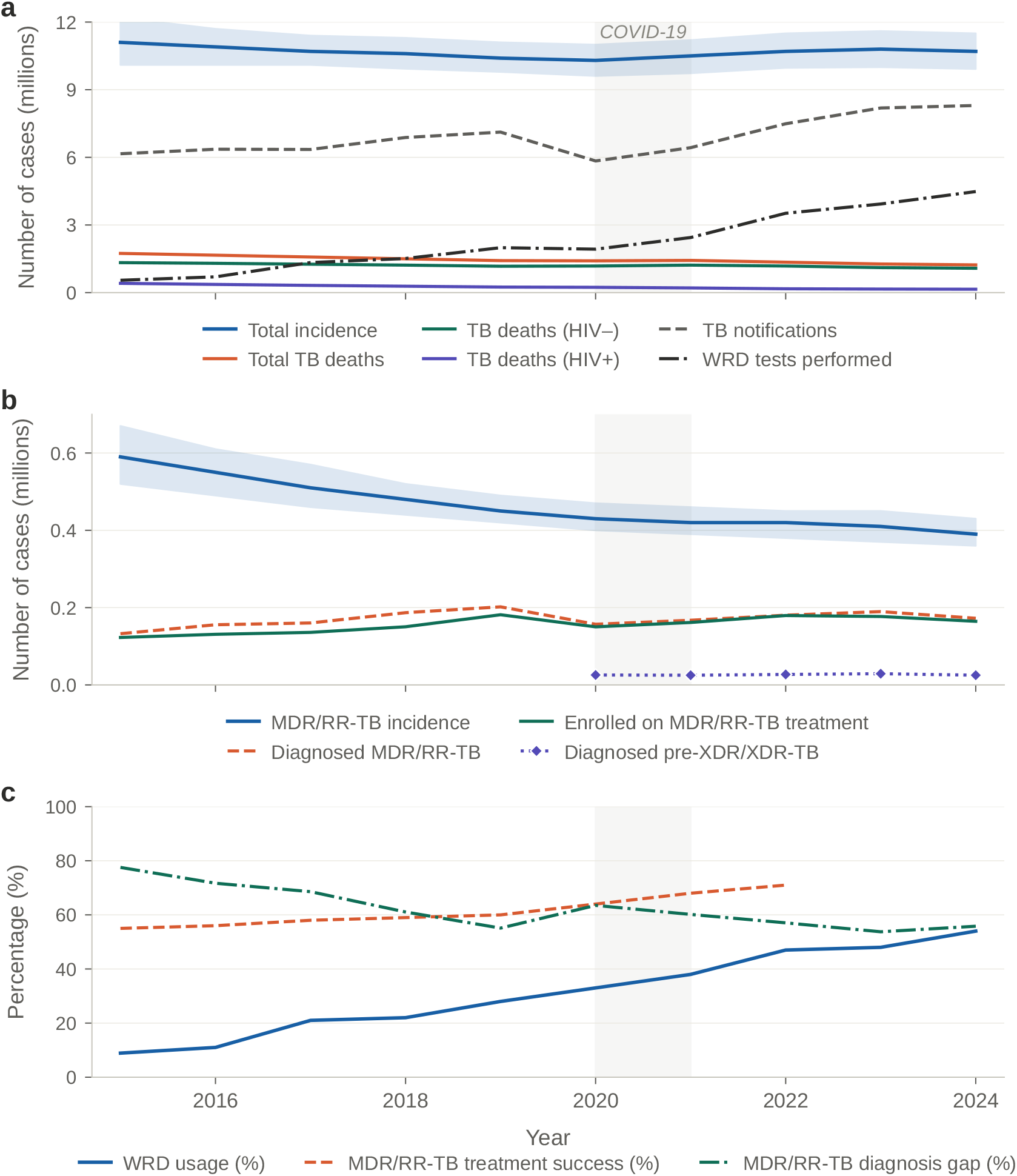
Global tuberculosis burden and programmatic indicators, 2015–2024. **a**, TB incidence, mortality (by HIV status), notifications, and WRD tests performed. **b**, MDR/RR-TB incidence, diagnosis, and treatment enrolment. **c**, WRD usage, MDR/RR-TB treatment success, and diagnosis gap. Shaded areas: 95% uncertainty intervals; grey band: COVID-19 period (2020– 2021). Data: WHO Global TB Report 2025.

**Figure 2:**
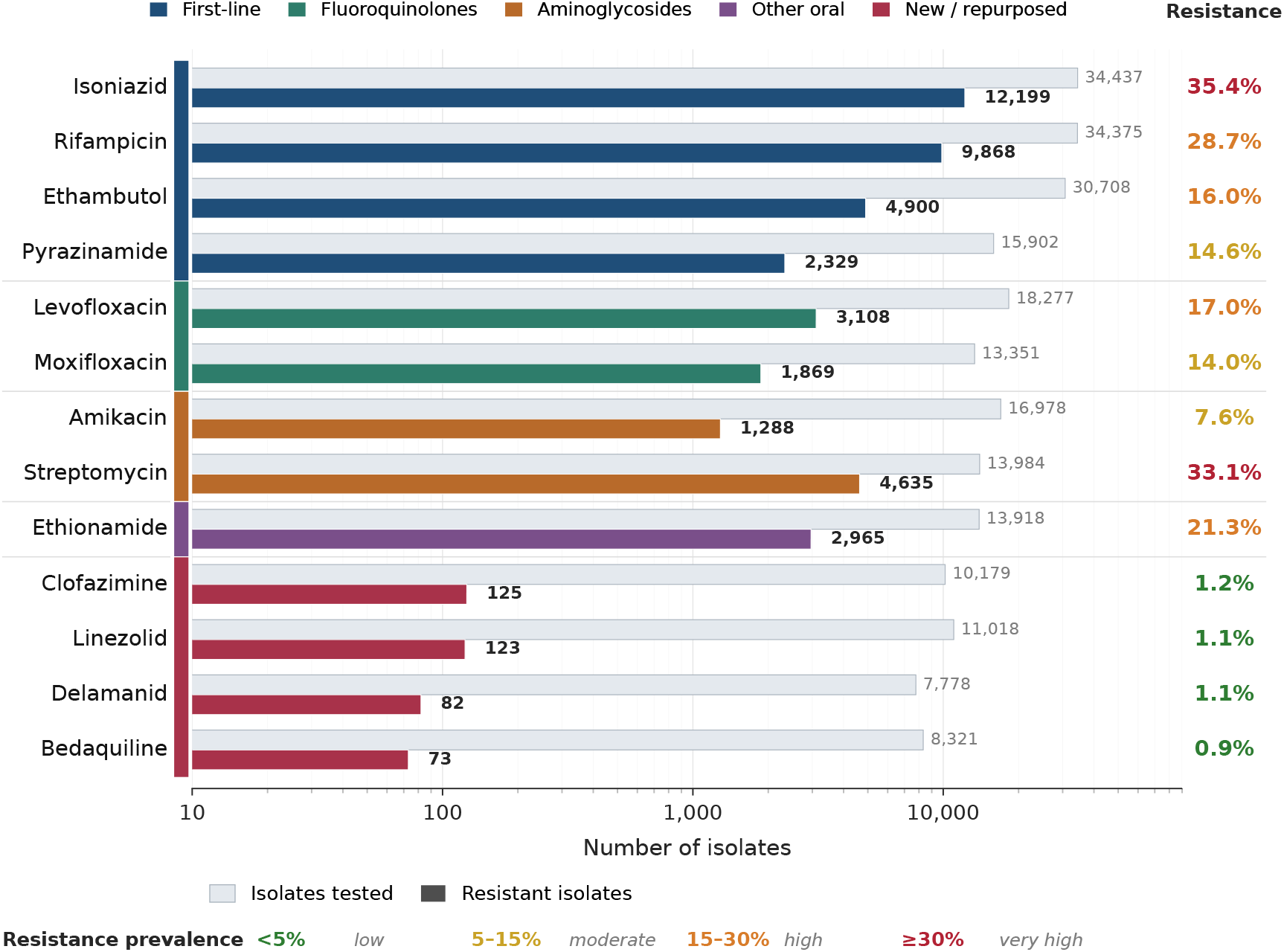
Phenotypic antibiotic susceptibility testing across 13 antimicrobials. For each antibiotic, the light grey bar shows the number of MTBC complex isolates with a phenotypic result and the coloured bar the subset classified as resistant. Resistance prevalence is calculated as *R/*(*R* + *S*). Data are from Table 1 of Walker et al. [49], the largest globally sourced MTBC dataset assembled to date for drug-resistance analysis including 38,215 isolates from 45 countries.

This review synthesizes the origin, phylogeny, and global distribution of MTBC lineages; the current understanding of drug-resistance mechanisms in MTB. We then examine computational approaches for predicting antimicrobial resistance, comparing catalog-based and *de novo* methods across their assumptions, data requirements, modeling strategies, and drug coverage, and *de novo* models where complex or previously uncharacterized mechanisms can be captured. We also address key barriers to deployment, including dataset bias, phenotype uncertainty, and limited generalization across populations. The goal is to clarify the state of the field, identify gaps that hinder clinical translation, and outline priorities for robust, scalable, and clinically actionable resistance-prediction tools for TB care.

**Table 1:**
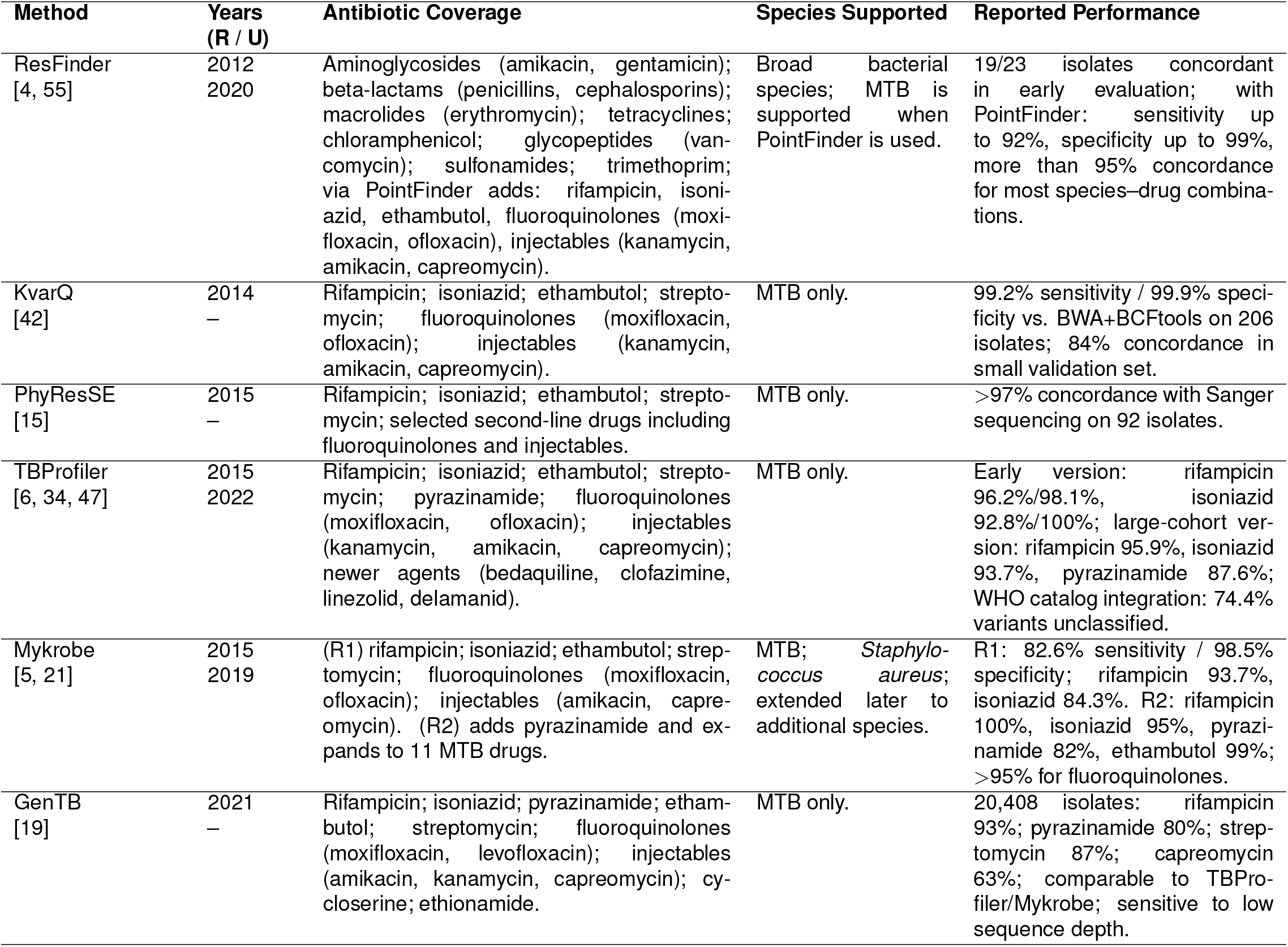
Catalog-based AMR prediction tools: antibiotic coverage, supported species, and reported performance. R = first release; U = last update.

## 2 Origin, Phylogeny, and Ecology of Mycobacterium

Understanding the evolutionary history and population structure of the MTBC is essential for interpreting patterns of drug resistance and informing lineage-aware control strategies. The MTBC comprises a clonal lineage of closely related bacteria that evolved from an environmental *Mycobacterium* ancestor into an obligate pathogen through step-wise adaptation to an intracellular milieu, a process hypothesised to have involved transitions from survival in free-living protozoa to replication in mammalian macrophages and, ultimately, to direct airborne transmission between human hosts [17]. Comparative genomics indicates that this transition involved genome downsizing relative to non-tuberculous mycobacteria coupled with horizontal gene transfer, although no single genomic feature is sufficient to account for the full virulence and transmissibility of the MTBC [17]. The closest known relatives of the MTBC are *M. canettii* and other smooth tubercle bacilli, all of which are strongly associated with the Horn of Africa; this geographic restriction, together with the concentration of human-adapted lineage diversity on the African continent, supports an African origin for the complex. The age of the most recent common ancestor remains unresolved, with ancient-DNA-based calibrations yielding estimates of less than 6,000 years, whereas models based on co-divergence with *Homo sapiens* suggest an origin approximately 70,000 years ago [17, 18].

The clonality of MTBC and its phylogenetic structure arise from the absence of ongoing horizontal gene transfer; diversification proceeds exclusively through single-nucleotide polymorphisms and chromosomal deletions, insertions, duplications, conversions, and inversions [18]. Within this framework, the number of recognised lineages has expanded over time. Gagneux [17] in 2018 described seven human-adapted lineages (L1–L7), whereas Goig et al. [18] in 2025 recognise ten human-adapted lineages (L1–L10), nine animal-adapted lineages, and one extinct lineage recovered from ancient human remains; Warner et al. [50] similarly refer to ten human-adapted and nine animal-adapted lineages. The phylogeographic structure of these lineages is marked by a contrast between globally distributed and geographically restricted variants: L4 occurs worldwide; L1 and L3 predominate around the Indian Ocean; L2 dominates in East Asia; L5 and L6 are largely confined to West Africa; L7 is almost exclusive to Ethiopia; and the recently described L8, L9, and L10 are each restricted to specific regions of Africa [18] (see Figure 3).

**Figure 3:**
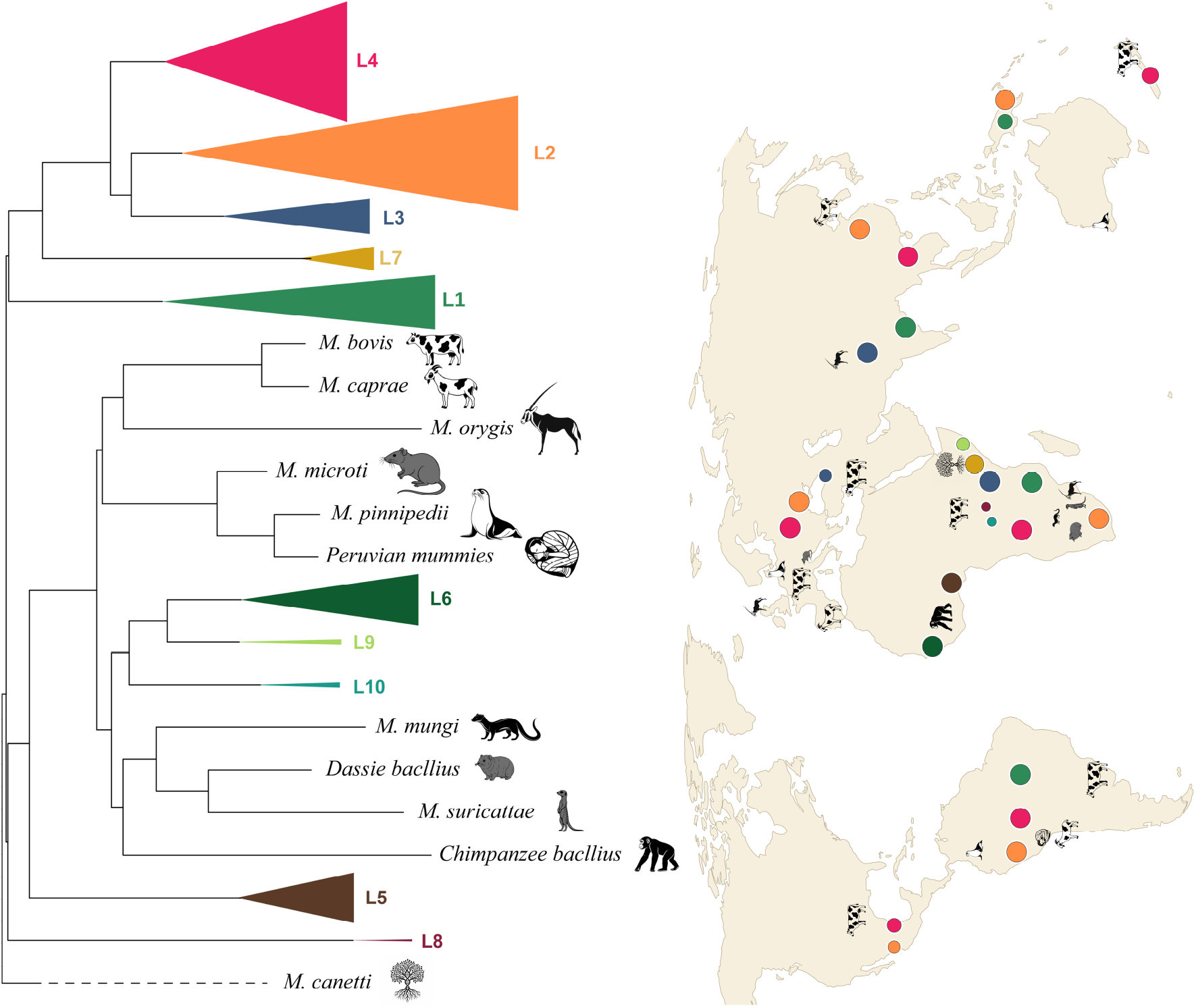
Phylogeny and global distribution of MTBC. The phylogenetic tree of the MTBC, rooted with M. canettii, includes 10 human-adapted lineages (shown as colored triangles) along-side multiple animal-adapted lineages. The leaves in the same lineage are collapsed for clarity. The map illustrates the worldwide geographic distribution of the MTBC lineages, with colored dots corresponding to their respective human-adapted lineages in the phylogeny. Following the pipeline from [60], we expanded the dataset to include additional genomes and processed all samples through BWA alignment [25] and bcftools variant calling [9] to create a comprehensive SNP alignment matrix for RAxML phylogenetic reconstruction [24], with detailed methodology and sample information provided in the supplementary materials.

Ecologically, this phylogeographic structure translates into a distinction between specialist and generalist lineages [18], a diversity that directly shapes AMR. Strict clonality means all resistance arises chromosomally, with its fitness cost modulated by the strain’s phylogenetic background and compensatory mutations [17]. L2 is disproportionately associated with MDR-TB, in part because multidrug resistance (MDR) L2 strains can acquire compensatory mutations that restore transmission fitness to the level of drug-susceptible strains, whereas MDR L4 strains retain a fitness cost that limits their spread. Goig et al. [18] frame this as historical contingency: lineages previously selected for virulence may have been predisposed to become dominant MDR genotypes once drug pressure was applied. Lineage-specific variation in intrinsic susceptibility—such as elevated pretomanid minimum inhibitory concentration (MIC) values in L1 [18, 50]—further ties the geographic distribution of lineages to the regional effectiveness of new treatment regimens. Surveillance and treatment strategies must therefore account for the lineage composition of local MTBC populations.

## 3 Mechanisms of Resistance and Their Drivers in Tuberculosis

In MTB, clinical drug resistance arises predominantly through *de novo* chromosomal mutations that alter drug targets, impair prodrug activation, or otherwise reduce drug efficacy [20, 31, 45, 57]. Altered efflux pump activity can also contribute, though current evidence suggests that, in MTB, efflux more commonly serves as an early adaptive *stepping stone* that permits bacterial survival at higher drug concentrations, rather than as the principal basis of clinically significant multidrug resistance [20, 45]. Higher-level resistance is instead driven by the sequential acquisition and accumulation of additional resistance-conferring chromosomal mutations [20, 31, 45, 57].This distinguishes MTB from many other bacterial pathogens, in which AMR commonly emerges through two broad genetic routes: mutation of chromosomal genes, and the acquisition of resistance determinants via horizontal gene transfer [28, 38, 48]. Horizontal gene transfer enables bacteria to acquire resistance genes through transformation, transduction, and conjugation [28, 48]. These processes are frequently mediated by mobile genetic elements, including plasmids and transposons, while integrons, often embedded within such elements, can capture resistance gene cassettes and provide the machinery required for their expression [28, 38, 48]. In many bacterial pathogens, such mobile elements are major drivers of multidrug resistance, facilitating both the acquisition and dissemination of multiple resistance determinants within and between bacterial populations [38, 48]. By contrast, current evidence indicates that horizontally acquired resistance genes play no major role in resistance in MTB [20, 31, 45, 57].

The emergence of drug-resistant TB is shaped not only by the molecular basis of resistance but also by treatment and transmission dynamics. The interplay between exposure to anti-TB drugs during treatment, person-to-person transmission, global travel, and poor-quality TB care has contributed to the emergence and establishment of distinct drug-resistant MTB strains in geographically distinct regions across the globe [13]. The standard 6-month regimen for drug-susceptible TB is driven by the duration required to cure the most severe forms of pulmonary disease, while shorter treatment durations appear sufficient for patients with less extensive disease [10, 12]; patient-level analyses show that the one-size-fits-all model leads to undertreatment of patients with severe disease and unnecessarily prolonged therapy for those with lower disease burden [22]. Therefore, personalized treatment tailored to the drug susceptibility profile of the infecting strain is recommended, and TB drug resistance detection is central to enabling such individualized decisions while minimizing selective pressures that drive further resistance acquisition and transmission [13, 45], with in vitro drug susceptibility testing being the recommended approach described in the following section [13, 45].

## 4 In Silico Identification of Antibiotic Resistance

### 4.1 Catalog-based methods

Catalog-based methods predict AMR in MTB by comparing WGS data from an isolate against curated databases of resistance-confering variants. Sequencing reads are usually aligned to a reference genome, single nucleotide polymorphisms (SNPs) and small indels are called at known resistance-conferring loci, and the resulting variants are matched against catalog entries to infer a resistance phenotype. Tools in this class include ResFinder [4, 55], KvarQ [42], PhyResSE [15], TBProfiler [6, 34, 47], Mykrobe [5, 21], and GenTB [19]. They differ in catalog composition, alignment and variant-calling strategy, antibiotic coverage, and the extent to which machine learning is integrated into the prediction step. Table 1 summarizes these methods; per-tool descriptions of the underlying database, workflow, and reported performance are given in the supplementary material.

### 4.2 *De novo* methods

*De novo* methods predict AMR in MTB directly from WGS data, without a predefined list of resistance-conferring variants. Reads or assembled contigs are decomposed into *k*-mers or gene-level features, statistical or machine learning techniques select features associated with resistance, and a predictive model maps these features to a phenotype. Because the feature space spans the entire genome, such methods can in principle capture resistance determinants that are absent from existing catalogs. Recent MTB-specific examples include the CRyPTIC machine learning models [8], MTB++ [40], and LLMTB [43], spanning gradient-boosted regression of MICs, sparse linear and ensemble classifiers, and transformer-based language models. LLMTB is more accurately described as a hybrid approach, as it restricts the input space to a curated set of resistance-associated loci but learns resistance signatures from sequence rather than matching against a predefined variant list. These methods differ in input representation, feature selection, and model architecture. Table 2 summarizes them; per-tool descriptions of training data, workflow, and reported performance are given in the supplementary material.

**Table 2:** *De novo* AMR prediction methods for MTB: antibiotic coverage, model architecture, training data, and reported performance. R = first release; U = last update.

| Method | Years<br>(R / U) | Antibiotic Coverage | Model Architecture | Training Data | Reported Performance |
| --- | --- | --- | --- | --- | --- |
| CRyPTIC ML<br>[8] | 2024<br>– | First-line (isoniazid, rifampicin, ethambutol); second-line (rifabutin, moxifloxacin, levofloxacin, amikacin, kanamycin, ethionamide); newer/repurposed (bedaquiline, delamanid, linezolid, clofazimine). | XGBoost regression on 31-mer features selected by univariate F-test; predicts $\log_2$ MIC, converted to binary calls via Youden's $J$. | 10,859 MTB isolates from the CRyPTIC dataset [7] (75% training / 25% testing). | External validation on 15,239 Seq&Treat isolates: sensitivity >90% for all drugs except ethionamide, clofazimine, linezolid; specificity >95% for all except ethambutol, ethionamide, bedaquiline, delamanid, clofazimine. |
| MTB++<br>[40] | 2024<br>– | 13 antibiotics: first-line (isoniazid, rifampicin, ethambutol); second-line (rifabutin, moxifloxacin, levofloxacin, amikacin, kanamycin, ethionamide); newer/repurposed (bedaquiline, delamanid, linezolid, clofazimine). | Dual classifiers per drug: least absolute shrinkage and selection operator logistic regression and random forest (RF) on 31-mer features selected by univariate $\chi^2$ test; 26 models total. | 6,224 MTB isolates from the CRyPTIC dataset [7] (80% training / 20% testing); five-fold cross-validation. | F1 >80% for first-line drugs; F1 >75% for most antibiotics; outperformed TBProfiler, ResFinder, KvarQ, and Mykrobe on most drugs, with gains driven by higher precision. |
| LLMTB<br>[43] | 2025<br>– | 13 antibiotics: first-line (isoniazid, rifampicin, ethambutol); second-line (rifabutin, moxifloxacin, levofloxacin, amikacin, kanamycin, ethionamide); newer/repurposed (bedaquiline, delamanid, linezolid, clofazimine). | BERT-inspired transformer encoding 273 curated resistance-associated genes (with 350 bp flanking regions) as single tokens; trained with focal loss, adversarial adjustment, and cosine annealing. | 12,204 MTB isolates from the CRyPTIC dataset [7] (6,250 development / 5,954 independent test). | Competitive F1-scores vs. TBProfiler, ResFinder, MTB++, and Mykrobe; attention patterns align with known resistance determinants (*rpoB*, *katG*, *inhA*, *rrs*, *gyrA/gyrB*, *embB*); reduced performance for linezolid, clofazimine, delamanid, bedaquiline. |

## 5 Discussion

TB remains a leading cause of infectious disease mortality worldwide, and the expansion of drug-resistant strains continues to erode treatment outcomes [54]. Although antibiotic susceptibility testing (AST) remains the gold standard for guiding treatment decisions, for MTB it routinely takes 6–8 weeks [58]. WGS-based prediction can return resistance calls in seconds to minutes per genome once sequencing is complete [16, 40]. Current WHO guidelines [51] accordingly endorse rapid resistance profiling to guide the initiation or adjustment of therapy pending AST results, shortening the interval to effective treatment without displacing the established clinical standard. This survey examines genomic methods of prediction of antibiotic resistance, focusing on their concordance with culture-based AST and their suitability for use within this pre-AST decision window.

WGS-based resistance predictors fall into two broad families: (i) catalog-based tools grounded in curated variant lists [16, 21, 47], and (ii) *de novo* approaches that learn directly from sequence-level features [8, 40]. The two families differ in how computational and curation costs are distributed across development and deployment. Catalog-based methods impose minimal computational demands at deployment, though this efficiency depends on substantial upstream curation [49] aggregating variant-level evidence from laboratories and consortia worldwide. Their principal constraints are scope and update latency: mechanisms absent from the reference list are missed however prevalent, and the lag between variant discovery and catalog incorporation imposes a structural delay, particularly consequential for emerging resistance mechanisms in newer drugs.

Addressing these blind spots of catalog-based methods motivates *de novo* frameworks such as MTB++ [40] and CRyPTIC-machine-learning [8], which capture multivariate and noncanonical resistance signals beyond catalog boundaries; aligning their significant *k*-mers to a reference genome further exposes locus-level associations. These gains come at substantial cost: MTB++ training requires hundreds of gigabytes of memory [40], and CRyPTIC-scale analyses require multi-terabyte memory distributed across multiple computational nodes [8]. *De novo* methods uncover associations that are also bounded by the genetic diversity of the training data, leaving mechanisms specific to underrepresented lineages undetected. The training data also constrain these methods in another way: because current *de novo* frameworks are developed on short-read sequencing, the distinct error and coverage profiles of long-read data mean that models trained on one modality may not generalize to the other without recalibration. Phenotypic label heterogeneity enhances the limitations of *de novo* methods: consortia such as CRyPTIC [7] adopt their own epidemiological cut-offs, so identical MIC values can yield discordant labels across repositories, a problem that standardized reporting of raw MIC values alongside binary classifications would mitigate.

One way to sidestep the label-heterogeneity problem is to predict MIC values directly rather than binary phenotypes. This has been demonstrated by the CRyPTIC consortium [8], whose models predict log_2_(MIC) and translate the values into binary resistance calls; however, the trained models were not released, and to our knowledge no other pre-trained tool provides this capability, nor does any comparable MIC dataset exist outside the CRyPTIC dataset [7]. Prediction of MIC offers advantages over binary classification: it preserves the resistance gradient that binarization discards, mitigates class-imbalance problems for antibiotics with few resistant isolates, and could inform dose optimization. Dose optimization matters because concentrations below the MIC allow resistant sub-populations to expand, while concentrations well above it increase the risk of drug toxicity.

Beyond data representation, the choice of model architecture also merits consideration, as large language models do not invariably outperform classical approaches. MTB is an instructive case: the species is highly clonal [17] and resistance is driven by a small, well-characterized set of genes, which limits the representational capacity that transformer architectures can productively exploit. Testagrose et al. [43] evaluated LLMTB against MTB++ [40] on a shared dataset and reported that MTB++ can matched or exceeded LLMTB on the antibiotics tested. On rifampicin, MTB++ achieved 89.86% accuracy against LLMTB’s 90.94%; on isoniazid, MTB++ reached 94.90% against LLMTB’s 93.61%. LLMTB, moreover, is not a *de novo* method. It relies on loci previously reported by CRyPTIC [7] and MTB++ [40], whereas the random forest (RF) and logistic regression (LR) classifiers in MTB++ are trained fully *de novo* from *k*-mer features. Once feature selection identifies the informative *k*-mers, RF and LR match or exceed transformer-based architectures at a fraction of the computational cost, an advantage that will become more pronounced as surveillance scales toward millions of isolates.

Rifampicin and isoniazid illustrate this discovery pathway in practice. For rifampicin, structural analysis of *rpoB* hotspot mutations shows that many reduce drug binding by disrupting hydrogen-bonding and van der Waals contacts with RpoB [26], motivating alternative RNA polymerase inhibitors with limited rifamycin cross-resistance [3, 23]. Isoniazid resistance commonly arises through reduced KatG-mediated activation or mutations in the *inhA* coding or promoter region, which has driven the development of direct InhA inhibitors that bypass KatG [3, 39]; candidate scaffolds include hydroxy-pyridones, thiadiazoles, diazaborines, and pyridomycin, several of which retain activity against isoniazid-resistant strains [3, 27, 41].

Pyrazinamide presents a different challenge, a drug whose clinical importance exceeds the reliability of current resistance testing. It occupies a distinctive role among first-line antitubercular agents owing to its sterilizing activity against persistent, slowly metabolizing bacilli in acidic microenvironments such as macrophages and caseous lesions [29], a property that, despite negligible early bactericidal activity, enables the six-month modern regimen [29]. However, phenotypic *in vitro* pyrazinamide AST of MTB lacks a true gold standard, has slow culture-based turnaround, and requires an acidic pH that itself inhibits MTB growth, so small variations in pH or inoculum size produce large MIC shifts and poor reproducibility [35, 46]. Genotypic alternatives based on *pncA* sequencing reach only 87% pooled sensitivity, as resistance-conferring mutations are scattered across the gene and additional mechanisms in *rpsA, panD*, and *clpC1* remain uncaptured by routine assays [2, 35, 49]. Among *in silico* tools, TBProfiler reached the highest sensitivity at 87.6%, with misclassifications attributed to rare *pncA* variants, nearly half unique to single isolates, absent from its catalog [34]. Pyrazinamide thus illustrates a case where, with rigorous quality control of phenotypic labels and training on diverse, high-quality strains, improved *in silico* prediction in clinical settings appears feasible.

The global burden of drug-resistant TB remains substantial: as shown in Figure 1b, annual detection of MDR-TB or RR-TB has exceeded 150,000 cases over the past five years, with pre-extensively drug-resistant TB (XDR-TB) or XDR-TB surpassing 25,000 each year [53]. Yet the largest MTBC dataset assembled for genotype–phenotype analysis, the WHO 2021 mutation catalog shown in Figure 2, contains at most 12,200 resistant isolates for any single antibiotic and under 130 for each of the new repurposed drugs (NRDs) central to MDR-TB/XDR-TB regimens: bedaquiline, delamanid, linezolid, and clofazimine. Molecular diagnostics and resistance catalogs for the most clinically important drugs are therefore built from a small fraction of circulating resistant strains. Because machine learning performance is sensitive to training-set size in this regime, with predictive performance rising sharply once resistant isolates per drug exceed approximately 500 [40], even modest expansions of the resistant-isolate pool for the NRDs could yield substantial gains in predictive accuracy. Closing this data gap through coordinated phenotyping and WGS of clinically resistant isolates from high-burden settings is a priority if molecular diagnostics are to keep pace with the evolving burden of drug-resistant TB.

Figure 4 integrates the elements discussed above into a single translational pipeline. Paired WGS and phenotypic AST of patient isolates train continuously updated machine learning models that identify novel resistance-associated loci, whose causal role is then tested through functional assays and gene knockout studies. Confirmed causal variants support four downstream applications: antibiotic discovery, updated resistance catalogs, refined molecular diagnostic panels, and retraining of the *de novo* models. In parallel, model predictions undergo concordance benchmarking against phenotypic AST, with clinical deployment limited to models meeting endorsement thresholds set by the WHO, the US Food and Drug Administration (FDA), or the US Centers for Disease Control and Prevention (CDC); endorsed models then inform early empirical therapy, shortening the interval to regimen optimization in suspected drug-resistant TB. Treatment outcomes subsequently re-enter the training pool, closing a learning loop that progressively expands coverage of NRDs, refines diagnostic accuracy, and supports individualized treatment regimens in high-burden settings.

**Figure 4:**
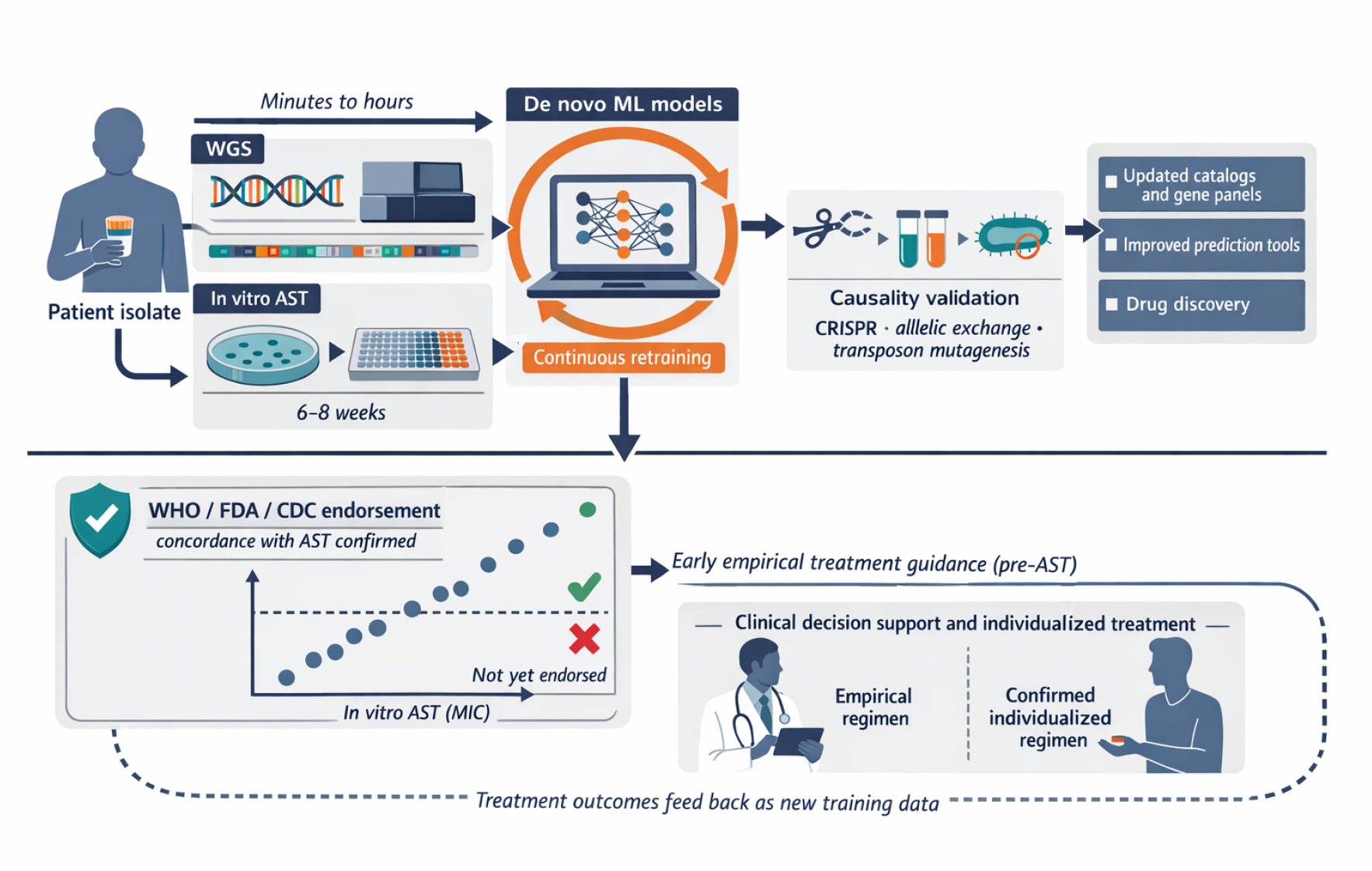
Translational workflow for WGS-based resistance prediction in TB. Patient isolates are processed in parallel through WGS and in vitro AST. Genomic and phenotypic data feed *de novo* machine learning models that identify novel resistance-associated loci, which undergo causality validation to produce updated catalogs, improved prediction tools, and drug discovery leads. These outputs inform clinical decision support and individualized treatment, with outcomes feeding back as new training data (dashed arrow).

## 6 Conclusion

In-silico prediction of antibiotic resistance in MTB offers practical decision support during the weeks-long wait for AST results, which can extend to months in resource-limited settings. Catalog-based tools such as TBProfiler [47], Mykrobe [21], and GenTB [19] deliver rapid, interpretable calls for well-characterized mechanisms, while *de novo* machine learning models [8, 40] surface additional resistance-associated loci and generate testable hypotheses about novel mechanisms. A central priority emerging from this work is quantitative phenotype modeling: predicting log_2_(MIC) values with calibrated uncertainty and deriving categorical labels only at the reporting stage, since MIC-level predictions accommodate site-specific breakpoints and enable individualized dosing assessments. Realizing these benefits depends on standardized MIC protocols, strict phenotypic provenance, and rigorous evaluation using lineage-aware splits and external validation across diverse geographies. Coupled with routine attribution of influential features to specific genomic loci, in-silico AMR prediction can provide reliable early treatment guidance while advancing mechanistic understanding of resistance in the MTBC.

